# Slow Dynamics Differentially Determine the Robustness of Regular Pacemaking: Distinct Subpopulations of Midbrain Dopamine Neurons Illustrate the Principle

**DOI:** 10.64898/2026.08.10.743865

**Authors:** Christopher J Knowlton, Strahinja Stojanovic, Marle Jahnke, Jochen Roeper, Carmen C Canavier

## Abstract

Pacemaking neurons, often found in mammalian nervous systems, integrate their inputs differently than quiescent neurons. Rhythmic single-spike pacemaking that is robust to noise can be achieved with a slow process that enforces a “resting potential” at each point along a ramp-like interspike interval (ISI) coupled with a fast restorative component. To demonstrate this phenomenon, we modeled previously identified distinct subpopulations of midbrain dopamine neurons that differed in projection target and in the regularity of their pacemaking. In the model of the more regularly-firing subpopulation projecting to the dorsomedial striatum, K_V_4 current was recruited by a deep after-hyperpolarizing potential (AHP) mediated by the SK channel. In the model of the less regularly-firing subpopulation projecting to the medial shell of the nucleus accumbens, the AHP was too shallow to recruit the K_V_4 current. In the more regularly firing population, the trajectory in the phase space of membrane potential and slow inactivation of K_V_4 was confined to move slowly through a narrow channel during the ramp-like portion of the ISI. Noisy perturbations from this channel were quickly damped by fast activation of K_V_4. In contrast, the smaller AHP in the model of the subpopulation projecting to the medial shell of the nucleus accumbens failed to recruit Kv4-mediated current, therefore the narrow channel was never entered, greatly decreasing the regularity in the presence of noise. This mechanism may be broadly applicable to single-spike pacemakers and explains how slow pacemaking with small net currents can be robust to fluctuations in single channel openings.

**Author Summary:** Pacemaking cells spike at regular intervals without the need for external input. There are numerous examples of pacemaking cells in the nervous system. We show that a process with slow dynamics relative to the individual spikes can make regular pacemaking robust to the noise that is always present in biological systems.

## Introduction

Generally, a neuron is envisioned as a quiescent cell that integrates its inputs to decide when to fire an action potential. However, spontaneously pacemaking neurons are found in many brain areas, including ithe basal ganglia, the cerebellum, and the hypothalamus; the inhibitory principal cells in the cerebellar cortex (1), the deep cerebellar nuclei (2), globus pallidus and the substantia nigra pars reticulata, as well as the excitatory principal cells in the subthalamic nucleus all pace. Many neuromodulatory neurons are also pacemakers (3). Moreover, pacemaking cardiac cells are found in the sinoatrial node and the atrioventricular node. Pacemakers integrate their synaptic inputs differently than quiescent neurons (4) because inputs to a repetitively firing neuron need not insert or delete spikes from the ongoing pattern; instead, they may alter the timing of spikes that would have occurred anyway. The window of temporal summation in repetitively firing neurons is not constrained by the membrane time constant; inputs arriving at any time during an interspike interval may influence the timing of the next spike. The effectiveness of inputs in altering spike timing depends not only on their sign and magnitude, but also on their time of arrival during the interspike interval.

Midbrain dopamine neurons constitute another example of spontaneous pacemakers in the basal ganglia (5). An influential theory is that a major function of midbrain dopamine signals is to report prediction errors (6). A level of spontaneous firing may provide a baseline from which both negative and positive prediction errors can be signaled. The regularity of pacemaking varies considerably between subpopulations of these neurons, thus we have chosen to model these subpopulations to provide insight into how regular pacemaking can be maintained in the presence of the noise that is invariable present in biological systems. For example, 10% of the VTA dopamine (DA) neurons (7) and 21% of SNC neurons (8) recorded *in vivo* exhibited regular pacemaking activity, despite being embedded within active neural circuits.

Two major electrophysiological phenotypes of dopamine neurons have been classified based on the intrinsic responses of these subpopulations to excitation and other attributes (9). Early studies focused on a generally more lateral, “conventional” population of DA neurons with projection targets in the dorsal striatum and the lateral shell of the nucleus accumbens (Fig 1A). These spontaneously pacing neurons (Fig. 1C1 and C2) exhibit very regular pacemaking, a pronounced after hyperpolarization (AHP), tall and wide (∼3 ms) action potentials with large spiking currents. They possess a limited dynamic range because they go into depolarization block at sustained firing rates above ∼10 Hz and they exhibit a prominent voltage sag upon hyperpolarization. During these early studies, DA neurons were thought to be a homogeneous population incapable of firing faster than 10 Hz in response to injection of depolarizing current (10,11) without simultaneous NMDA receptor activation.

**Figure 1:**
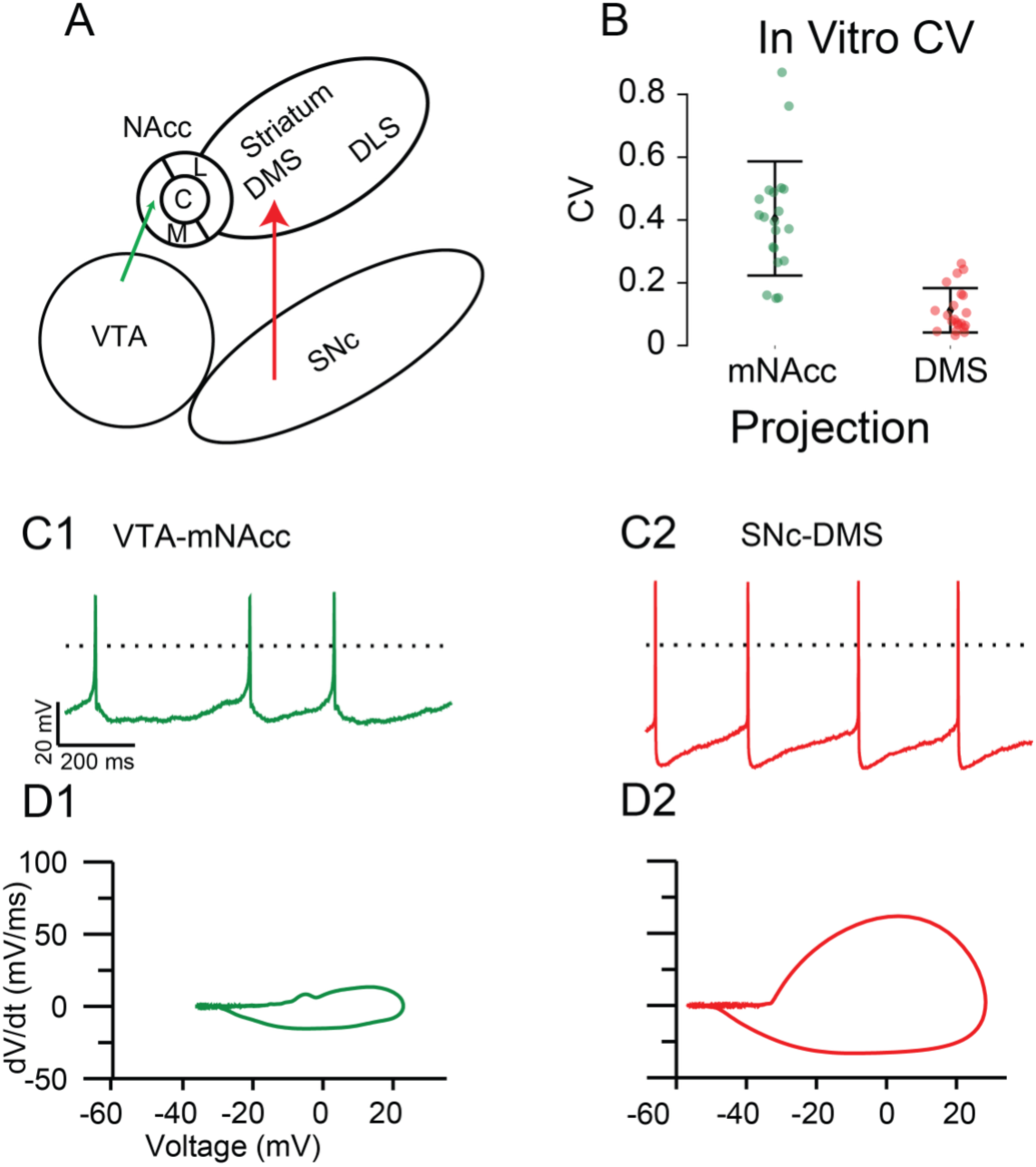
Subpopulation Electrophysiological Phenotypes with Exemplars. A: Schematic of a coronal slice of mouse midbrain (medial left, lateral right) with projection targets indicated by the arrow. Atypical subpopulation exemplars are the VTA-mNAcc (green) and SNc – DMS (red). B. Coefficients of variation for the various subpopulations in vitro. C. Representative Experimental Traces *in vitro* from the atypical subpopulation (C1) and the conventional subpopulation (C2). D. Phase plane plots of experimental traces above.

**Figure 2:**
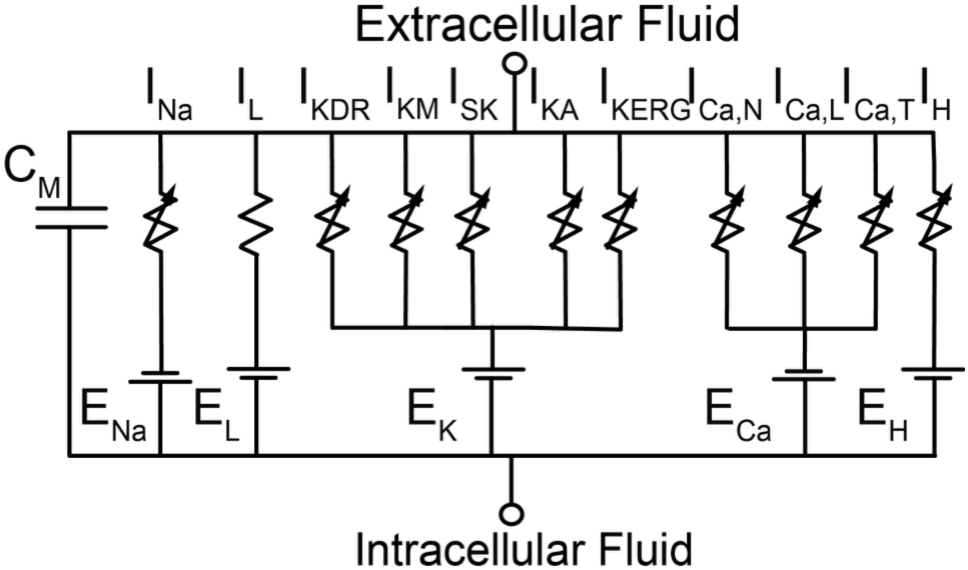
Model Schematic. Equivalent Circuit for Membrane Potential.

A second, “atypical” subpopulation was identified consisting of dopamine neurons projecting (Fig. 1A) to the prefrontal cortex, the medial shell and core of the nucleus accumbens, and other targets (9,12–15). These neurons possess an extended dynamic range, a less prominent AHP, little sag response, and smaller maximum current during spiking. Computational modeling (16) suggests that a lower availability of the FGF12 auxiliary subunit of the Na_V_ channel that promotes long term inactivation (17,18) enables the higher frequency in this atypical, extended dynamic range population, which can fire in the 12-20 Hz range in response to somatic depolarization in the absence of NMDA receptor activation (9).

To motivate the present study, we analyzed data from spontaneously active dopamine neurons identified by projection target, with representative examples shown in Fig 1C. Figure 1B shows that under control conditions *in vitro* (see Methods) atypical DA cells projecting to the mNAcc from the VTA have a larger coefficient of variation than the conventional ones projecting to DMS: mNAcc: 40.46+/-18.18% N=20, DMS: 11.22+/-7.07% N=21. (The conventional DLS projecting population is similar to the DMS projecting, see Supplementary Figure 1). In this study, we provide a mechanistic explanation for the differences in regularity.

For the specific example addressed in this study, defining the differential mechanisms that control DA subpopulation-specific intrinsic dynamics will not only provide a deeper understanding of their contribution to the behavioral dimension of dopamine in the context of motivated behavior but may also facilitate the identification of causal drivers in the pathophysiology of DA-related psychiatric (e.g. schizophrenia, depression, substance use disorders, ADHD) and neurological disorders (e.g. Parkinson Disease, Tourette Syndrome). However, the implications are much wider if, as we hypothesize, similar principles apply to other populations of pacemakers.The robustness of pacemaking determines the balance between maintaining a rhythmic firing rate and responding to perturbations. The more robust the regularity of pacemaking the less responsive the neuron is to external perturbations and vice versa.

### Mathematical and Computational Methods (detailed Computational and Experimental Methods follow the Discussion section)

The equivalent circuit model for our single compartment models is shown in Fig. 2. The models include a fast Na^+^ current (I_Na_) (16,19), a leak current (I_L_), and five K^+^ currents: a delayed rectifier (I_KDR_), the M-type K^+^ current (I_KM_) (20), a Ca^2+^ activated small conductance K^+^ current (I_SK_) (5), an A-type K+ current (I_KA_) mediated by K_V_4.3 (21), and the ether-a-gogo-related K+ current (I_KERG_) (19). There are also three Ca^2+^ currents, the N, L and T-type (_ICa,N_, I_Ca,L_, and I_Ca,T_) currents and a nonspecific cation H current (I_H_). Hodgkin and Huxley (22) type models were used for all current except I_Na_ and I_KERG_ for which Markov models were used. The most important inward currents for this study were I_Na_ and I_CaL_ and the most important outward currents were I_SK_ and I_KA_. The I_Na_ and I_CaL_ currents created the positive feedback that destabilized any resting potential and forced the neuron to pace.

In order to model I_SK_, a model of Ca^2+^ handling was implemented that included a microdomain (23) for the Ca^2+^ that activates that channel The Ca^2+^ sources that activate SK are not fully understood;however, the N and T-types are known to contribute, but not the L-type (24). Since we only use Ca^2+^ in this model to drive the SK channel, the parameters of the microdomain were adjusted to simulate the time course of the SK current and neglect the contribution of the L-type and part of the T-type channel to the Ca^2+^ dynamics. The parameters that differ between subpopulations are given in Table 1. The most important differences among the subpopulations were those in I_SK_, which controls the depth of the AHP, including the smaller conductance in the atypical population and the larger diameter parameter (dCa) that corresponds to a weaker coupling between the Ca^2+^ channels and SK (see detailed computational methods section). These differences affect the recruitment of I_KA_, whose time constant of inactivation is another important parameter that varies across subpopulations. The model can be accessed at https://modeldb.science/2041613 (reviewer password: nullcline).

**Table 1:** Key Model Parameters. Representative models have varied parameters based on projection target. The Kv4.3 with a slow auxiliary subunit is present only in VTA-mNAcc model. CI1I2 is the transition rate into the longterm inactivation of NaV. Full model equations are given in Supplemental Materials. dCa scales the effective size of the microdomain that activates the SK channel.

|  | VTA-mNAcc | SNC-DMS |
| --- | --- | --- |
| gNa ($\mu\text{S}/\text{cm}^2$) | 7000 | 15000 |
| gKDR ($\mu\text{S}/\text{cm}^2$) | 1400 | 2000 |
| tauKA (ms) | 44%: 65<br>56%: 150 | 75 |
| gCaT ($\mu\text{S}/\text{cm}^2$) | 0 | 10 |
| gCaN ($\mu\text{S}/\text{cm}^2$) | 200 | 200 |
| gCaL ($\mu\text{S}/\text{cm}^2$) | 5 | 10 |
| gSK ($\mu\text{S}/\text{cm}^2$) | 50 | 250 |
| gKA ($\mu\text{S}/\text{cm}^2$) | 187.5 | 468.75 |
| gKM ($\mu\text{S}/\text{cm}^2$) | 90 | 100 |
| gKERG ($\mu\text{S}/\text{cm}^2$) | 5 | 10 |
| gH ($\mu\text{S}/\text{cm}^2$) | 0 | 50 |
| gLNa ($\mu\text{S}/\text{cm}^2$) | 3 | 6 |
| gLK ($\mu\text{S}/\text{cm}^2$) | 5 | 9 |
| CI1I2 (/ms) | 0.05 | 0.2 |
| dCa ($\mu\text{m}$) | 0.2 | 0.1 |

#### Fast/slow phase plane analysis using nullclines

The intrinsic dynamics of pacemaking neurons can often best be understood using a technique called separation of time scales, in which the state variables (membrane potential, Hodgkin-Huxley type gating variables, concentration, and Markov model occupancy states) are subdivided into fast and slow groups based on their dynamics (25). We collapse the dynamics of all the fast gating variables by setting them to their steady value as a function of the membrane potential. This reduces the system to only two variables, membrane potential (V) and a much slower variable, which renders the system amenable to analysis in a phase plane (Fig. 3). The V nullcline (black curve) is the set of pairs of values of the membrane potential and the slow variable at which the net current (which determines the rate of change) is zero, and the slow nullcline (green curve) is the set of pairs of values at which the rate of change of the slow variable is zero. The only possible resting potential is at an intersection of the two curves. However, the V nullcline for a relaxation oscillator that supports pacemaking is nonmonotonic with three branches (26,27). The middle branch is created by the auto-catalytic, regenerative dynamics of currents like low threshold L-type Ca_V_1.3 and the persistent Na_V_, modeled as residual occupancy in the open state in our Markov model. Positive feedback from regenerative currents destabilizes the resting potential as any slight depolarization opens inward channels and recruits more channel opening, increasing the inward current. Conversely, any slight hyperpolarization closes channels and causes more inward channels to close. Therefore, trajectories are pushed away from this branch (arrows). Under the fast/slow separation of variables with the slow variable on the x-axis, movement in the vertical direction is fast and tends toward the stable branches of V nullcline and away from middle branch; the dynamics evolve slowly along the two stable V nullcline branches. The lower stable branch represents a hyperpolarized branch in which the channels mediating the regenerative subthreshold currents are closed, whereas on the upper branch these channels are open, but balanced by outward, restorative (where increasing voltage increases an hyperpolarizing rather than depolarizing) current. The direction of motion is determined by the slow variable, which decreases above and increases below the slow variable nullcline. Fig. 3 illustrates relaxation oscillator (28) dynamics in which the oscillatory trajectory (red curve) follows the fast (V) nullcline and jumps between branches at the bends often called knees.

**Figure 3:**
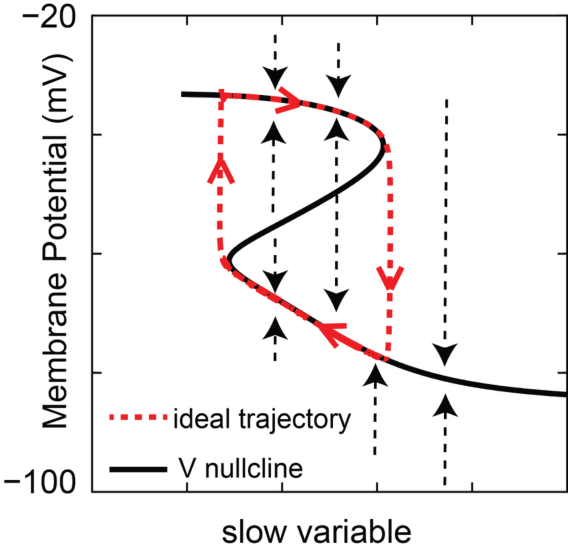
Fast/Slow analysis. The nullclines determine trajectories as shown by arrows denoting the vector field. Canavier et al., 2016

## Results

The mathematical phase plane analysis described above was applied to our biophysically calibrated models of the distinct subpopulations of dopamine neurons. First we studied the more regularly firing subpopulation. Fig 4A shows that the interspike interval of an identified DMS projecting DA cell has a ramplike shape in this subpopulation (29). The net current was calculated by scaling the derivative of the voltage by the capacitance of a typical SNc DA neuron (60 pF)(30). A low-pass filter (3^rd^ order Buttersworth with 500 Hz shoulder) and 50 Hz notch filter were applied to remove noise associated with known and likely artificial measurement noise sources. The ramp-like interspike intervals contain consistently small ∼1-2 pA mean net current as previously shown in (29). Fig 4B shows a representative voltage trace generated by a computational model using the channels from Figure 2 with a typical sequence of action potential (red), AHP (purple), and ramp (orange) and the corresponding transmembrane currents, calculated in the same way as in Fig. 4A but without the need for filtering. The AHP is driven primarily by SK currents recruited by high threshold Ca^2+^ channels during the preceding spike (24).

**Figure 4:**
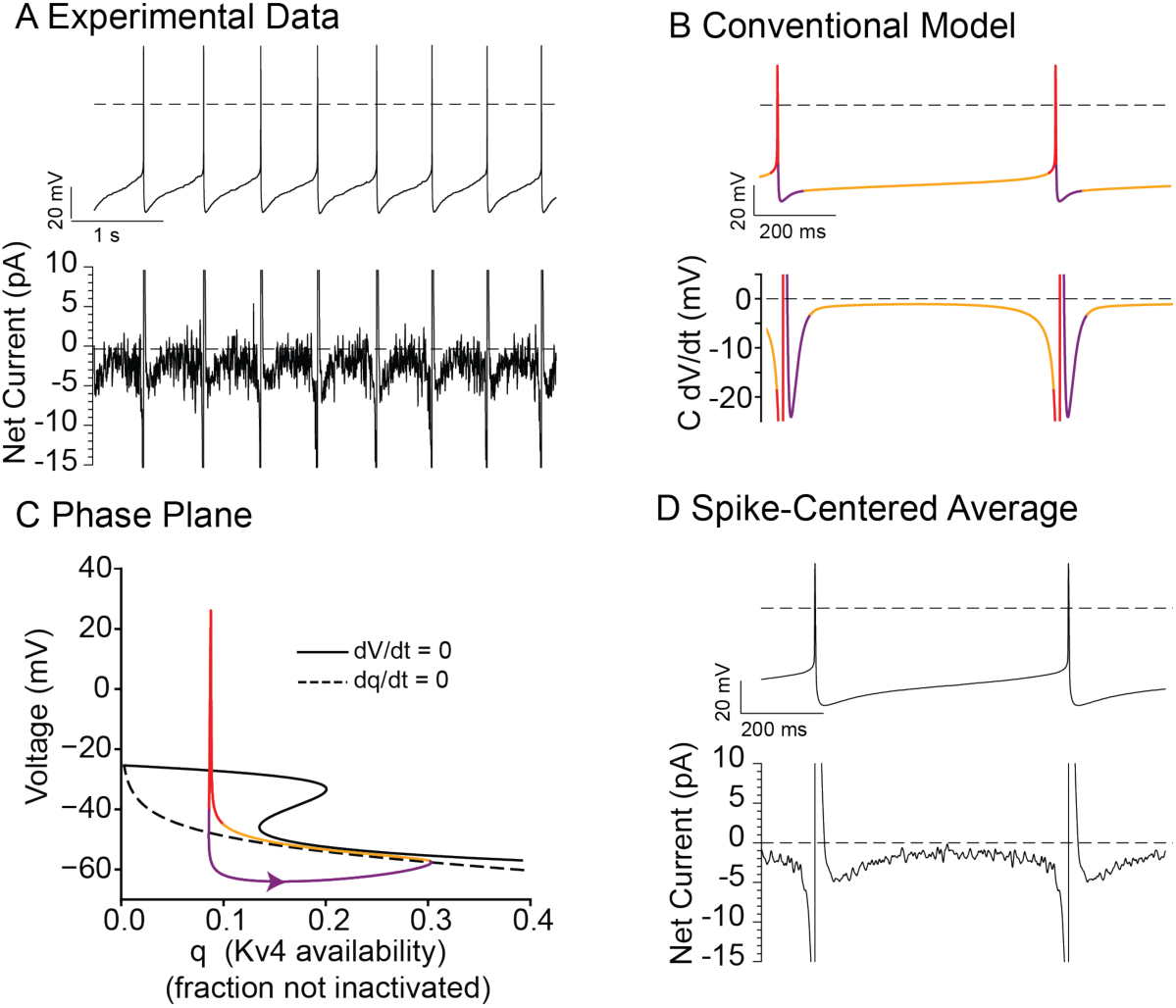
Fast/Slow analysis on SNc-DMS model. **A.** Experimental voltage trace (top) from a conventional neuron illustrating the ramp. Net current (bottom) estimated by filtering the voltage waveform with a500 Hz Buttersworth low-pass and 50 Hz notch filters and then multiplying −dV/dt by 60 pF.B: Voltage trace with color coded phases: action potential–red, AHP-purple, and ramp-orange. C: Phase space with respect to Kv4 inactivation. D. Spike-centered average of voltage, current traces in A (n=15 interior spikes). Pre, post spike waveforms are joined at −50 mV.

Figure 4C applies the concepts from Figure 3 to the model of a conventional DMS projecting DA neuron. Figure 4C shows the voltage nullcline with respect to the fraction of non-inactivated Kv4 ion channels required to produce sufficient outward A-type current to cancel out the steady state inward current at each voltage (black curve) and nullcline for the inactivation gate, which is the steady state non-inactivated fraction of Kv4 (dashed curve). The voltage nullcline is given by the pairs of points (V,q) that make the net current zero with all variables except q held to their steady state value. Additionally, the phase portrait of the dynamics is shown with colors corresponding to regions of the ISI from 4B in a clockwise direction. During the AHP indicated in purple, the dynamics are controlled by the slow dynamics of microdomain Ca^2+^ concentration, thus they are not constrained by the voltage nullcline (black curve) because that nullcline was calculated under the assumption that the slow inactivation of KV4 controls the dynamics. As the trajectory is now to the left of the steady state Kv4 inactivation at these voltages, the effect of the AHP is to move the trajectory to the right, towards the steady state inactivation curve, increasing the availability of the K_V_4 current by removing inactivation. The trajectory reverses direction and moves leftward only after crossing the Kv4 nullcline as the AHP currents deactivate. After the AHP and during the ramp-like part of the ISI, Fig. 4C shows that trajectory closely follows the lower, stable branch of the voltage nullcline, as expected if the inactivation of K_V_4 is the dominant slow variable since the voltage nullcline was calculated under that assumption. The steady state inactivation curve for the K_V_4 channel (dashed curve) is also a nullcline since the rate of change in the inactivation variable is zero on the curve (which would be a sigmoid if the axes were flipped). The trajectory is more precisely described as being confined to a narrow channel between the two nullclines. In this channel, the dynamics are dominated by the slow inactivation of the Kv4 ion channel, which causes a leftward movent on the yellow portion of the trajectory in Fig. 4C. Inactivation decreases outward current causing a slow upward shift along the same part of the trajectory. The reason the movement is slow is because that rate of inactivation is given by the following equation (see full model details in supporting information): *dq* / *dt* = (*q*_∞_ (*V*) − *q*) / τ*_q_*. In this equation, *q_∞_(V)* denotes the steady state value of the inactivation gating variable at a given membrane potential whereas *q* denotes the instantaneous value of the inactivation gate. The difference between these two values is small when the trajectory (q value) is near the steady state, and moreover the rate of change is obtained by dividing this small value by the relatively large (compared to the fast variables) inactivation time constant of ∼50 ms, making the dynamics slower than the ∼50 ms time constant of inactivation would suggest.

The trajectory is released from this channel to produce an action potential (red trace in Figure 4C) only after reaching the knee at the left end of the lower branch in Figure 4C. The dynamics during action potential (red trace) do not follow the upper branch of the voltage nullcline because that branch is not truly stable. Instead, it is a spiking branch, because it is above the threshold for action potential generation. Therefore, at least one more dimension orthogonal to the phase plane in Fig. 4C would technically be required to allow the trajectory to spiral around the top branch during spiking as in (19). Note that the q nullcline only controls the dynamics during the ISI and not during the other phases of the pacemaking cycle.

Figure 4D shows the effect of averaging across traces to remove intrinsic noise from the traces in 4A. A spike-centered mean pacemaker waveform was generated from n=15 recorded spikes and their surrounding interspike intervals from a single 10s in vitro recording of an identified DMS projecting SNc neuron. The full mean interspike was then generated by connecting the pre-AP and post-AP waveforms at their respective −50 mV crossings and normalizing to the average period. The resulting waveform matches that of the model in Fig. 4B, including an AHP with larger net current followed by an approximately constant net current ramp from ∼-55 mV until about −45 mV followed by the action potential.

In order to test the robustness of pacemaking in the above described exemplar from the conventional population, we applied a normally distributed, zero mean noise signal analogous to the intrinsic channel noise of recorded cells *in vitro*. Fig. 5A1 shows that in the conventional model of the DMS-projecting neurons, perturbations during the ramp portion of the ISI remain confined within the region between the nullclines (indicated by yellow oval), rapidly restoring the dynamics to the channel. Setting the SK conductance to zero to simulate the bath application of apamin reduces the AHP and largely prevents the removal of Kv4 inactivation. Since K_V_4 inactivation is not removed, the dynamics of the DMS projecting model no longer enter the channel that confines the trajectory and mitigates perturbations (Figure 5A2), thus identical input pertubations evoke larger voltage excursions that alter the timing of action potentials and decrease regularity. Fig. 5A3 illustrates the difference in CV caused by setting the SK conductance to zero.

**Figure 5:**
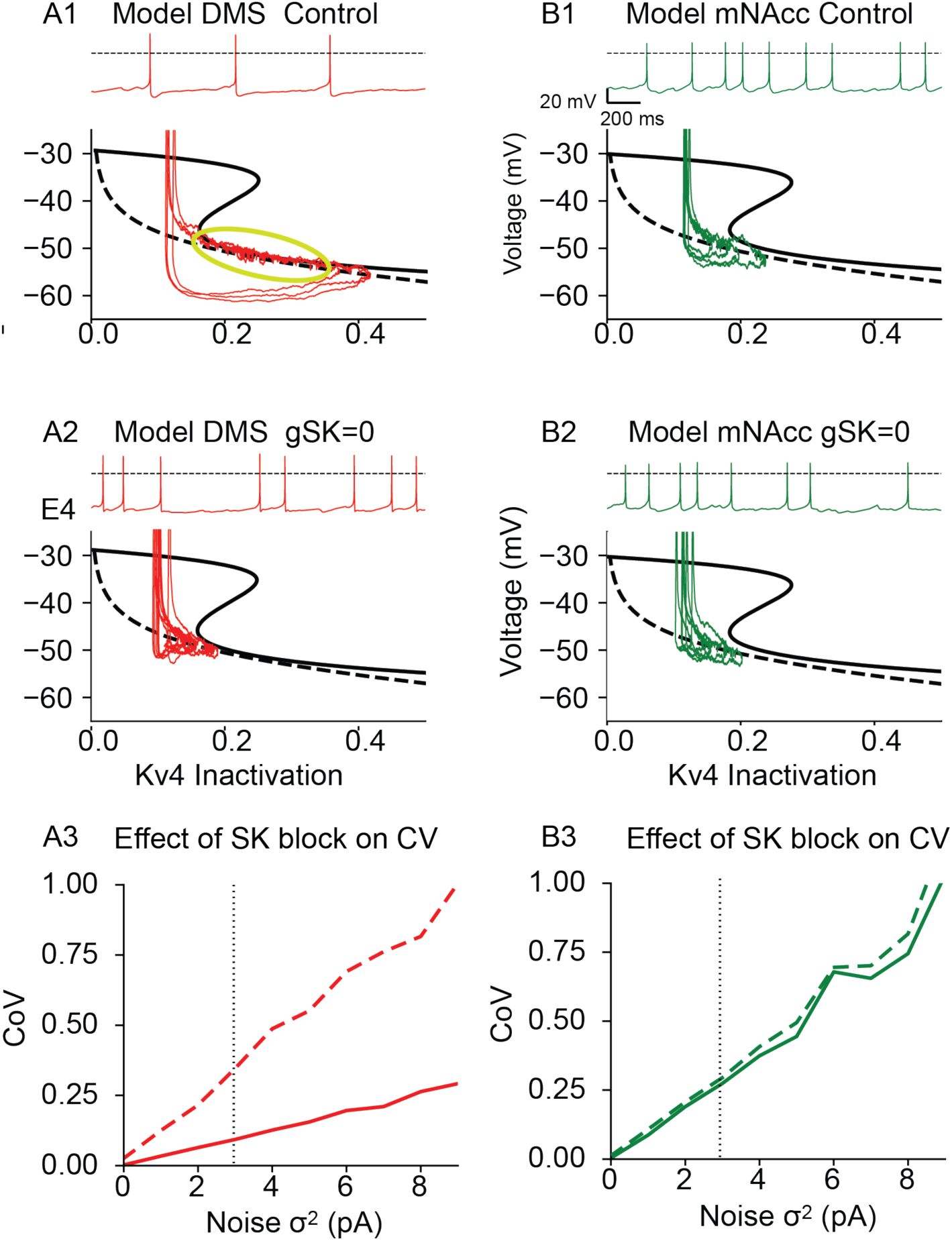
Narrow channel between nullclines constrains trajectory to promote regular ISIs. **A1**: top: Voltage trace of SNc-DMS model with 3 pA gaussian noise bottom: Phase plane representation. A2. Same as A1 but

We next investigated the pacemaking dynamics of the less regular atypical DA neurons represented by our model of the mNAcc projecting neurons. The single most reliable marker to discriminate between the conventional and atypical subpopulation pacemaker firing is that the atypical neurons consistently have a shallower AHP (12,27), mediated by a smaller SK conductance (putatively aided by weak coupling to fewer Ca^2+^ channels). This is consistent with the relative insensitivity to apamin in that population (31). Despite the presence of similar narrow channel between the Kv4 inactivation and voltage nullclines, the AHP is too shallow for substantial recruitment of Kv4 (Fig. 5B1), so the narrow channel is never reached. Therefore, perturbations in the mNAcc projecting model occur mostly in the unconstrained left section of phase space. In this region perturbations are not rapidly restored and can rapidly summate to produce rapid spiking. Moreover, if noise perturbs the trajectory into the narrow channel, the subsequent spike can be delayed. Since the AHP mediated by the SK channel is not sufficient to recruit K_V_4 and stabilize pacemaking, blocking SK in the atypical model has little effect on the trajectory (Fig. 5B2) or the CV (Fig. 5C3). Note that in the absence of the SK conductance the CV between the two populations is comparable.

We then used the electrophysiological concept of an instantaneous IV curve (27) to explain the strong restorative forces occurring within, but not outside of, the narrow channel between the nulclines. We generated these instantaneous IV curves by holding values of the slowest variables constant at their values at the colored dots on the simulated voltage traces in Fig. 6A1 and B1. The variables that were held constant were inactivation of K_V_4 (IA) and T-type Ca^2+^ channels, activation of the SK and M channels, and the open fraction of ERG channels. Then we plotted the steady state current evoked at membrane potentials starting from a hold at the indicated point and stepping to −56 to +40mV given that set of fixed slow variables. This provides a snapshot of the I-V curve on a time scale that is very fast relative to the period of the regular pacemaker. The slope of the instantaneous IV curves in Fig. 6A2 and B2 is equal to the instantaneous restorative conductance. The steep slopes in Fig. 6A2 indicate a strong restorative response; any small depolarization from the dot results in a strong outward current whereas any small hyperpolarization results in a strong inward current. The colored dots on the trajectory (gray) are fixed points on a fast time scale because brief perturbations of the membrane potential in either direction are pushed back to those points by a restorative current proportional to the steep slope of the instantaneous IV curves. The phase plane in Fig. 6A3 shows that the ramp response is confined to the region between the voltage and Kv4 nullclines. The region between those nullclines is analogous to a stable fixed point, or resting potential, that moves with the slow variable. This ‘moving fixed point’ is only truly fixed at fixed values of the slow inactivation; since the value of the inactivation changes slowly, the “resting” potential moves slowly. The instantaneous fixed point loses its influence on the dynamics when the Kv4 channel is replaced as the dominant channel by recruitment of regeneratively inward CaL and NaV channels at depolarized voltages. On the other hand, for the atypical model in Fig. 6B1, the failure to recruit Kv4 results in relatively flat instantaneous I-V curves (Fig. 6B2) during the ISI with small slopes, resulting in only weakly restorative or nonexistent Kv4-mediated currents. Therefore there are no fixed points of a fast time scale in the phase plane in Fig. 6B3.

## Discussion

### Generalization to other neuromodulatory populations

Neurons that release modulatory neurotransmitters such as dopamine, serotonin, norepinephrine and histamine constitute a major subcategory of spontaneous tonically and rhythmically firing neurons. These neurons typically fire single action potentials at slow rates (0.5-5 Hz) and provide a continual release of the neuromodulator in the projection target areas (32). The regularity of pacemaking in the other populations of neuromodulatory cells may be governed by the same principles as we described here for dopaminergic neurons. The best analogy can be made with the hypothalamic histaminergic tuberomammillary neurons that fire regularly in vitro at 0.5-10 Hz. They also exhibit an A-type current, have a ramp response to a hyperpolarizing step and a ramplike ISI that follows an AHP (Fig 2 C in (33)), which removes inactivation of the A-type current (34). Both of the DA neuron subpopulations modeled in this study also exhibit a ramp response (Fig. S2) due to an A-type current. Another neuromodulatory population, the serotonergic neurons in the dorsal raphe, exhibited extraordinary rhythmicity and a relatively slow rate of discharge (∼1.3 Hz) in anesthetized rats (35). Early electrophysiological experiments carried out in brain slices revealed that these neurons spontaneously discharge with a slow (1–2 Hz), regular (clock-like) pattern. As in dopaminergic and histaminergic neurons, the ISI in serotonergic neurons consists of an AHP (∼6 mV in that study) followed by a gradual, ramp-like interspike depolarization (36). Yet another neuromodulatory population, the locus coeruleus noradrenergic neurons in the pons fire spontaneously in rats at 0.5-2 Hz in vivo and at 1-5 Hz in vitro. The ISI was once again composed of an AHP that decayed fairly rapidly followed by a ramp to threshold (Fig 12C in (37)). The calculated net current during the ramp (illustrated in Fig 6A of (38)) is relatively constant and less than 1 pA. The similarity between the noradrenergic and the conventional dopaminergic waveforms is striking.

**Figure 6:**
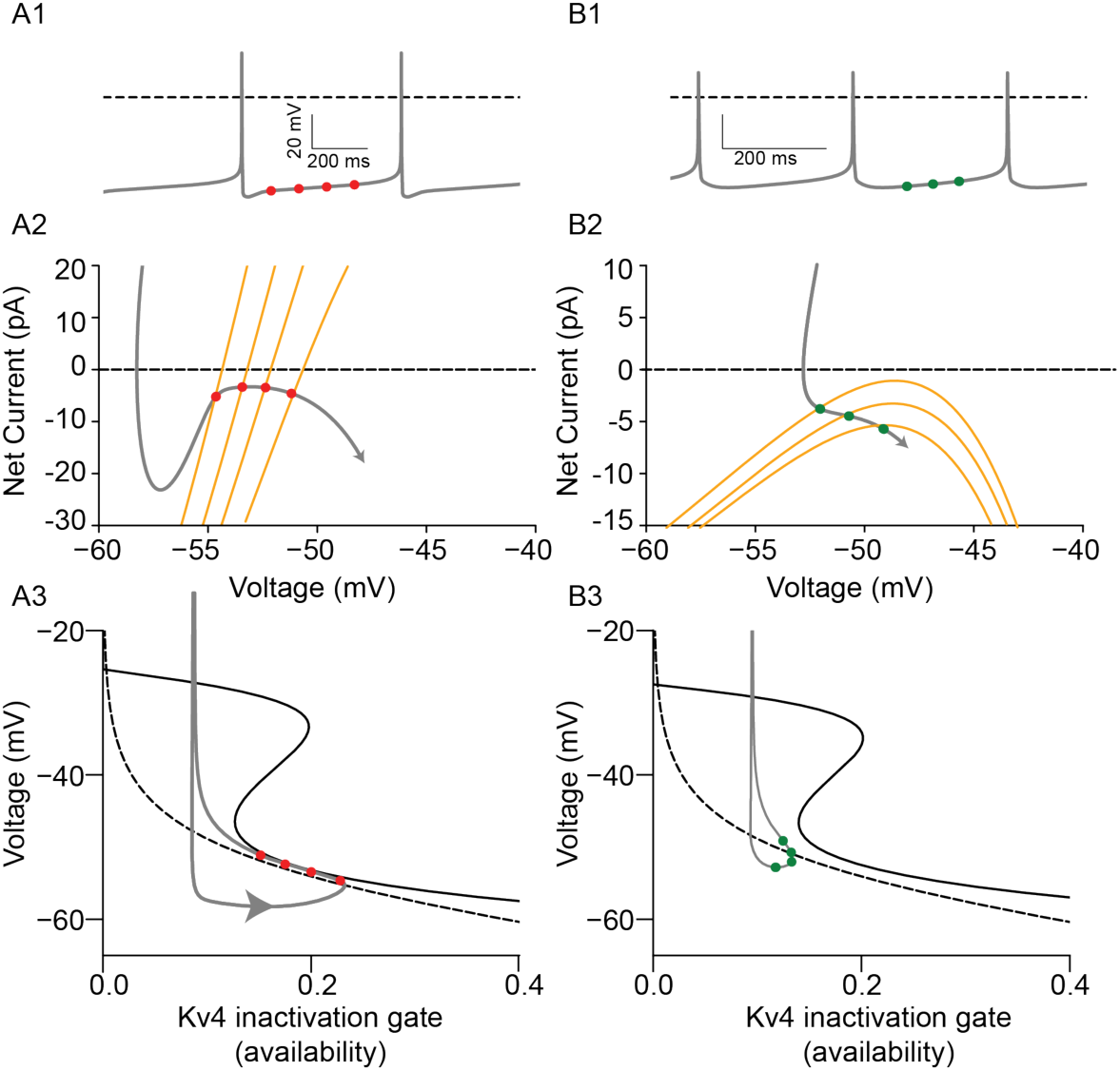
Instantaneous IV curves explain robustness. **A.** Conventional model neuron. A1. Voltage trace with sampled voltage points. A2. Instantaneous IV curves at each point (orange) and trajectory (gray). A3. Phase plane showing rolling fixed points (dots). **B**. Same as A for the atypical model.

### Robust slow pacemaking in the conventional population

An *in vitro* study of the regulation of pacemaking in VTA DA neurons focused selectively on the most regularly pacemaking neurons (29), which we now know were likely conventional neurons projecting to the lateral shell of the nucleus accumbens (9). That study showed that the interspike interval between two spikes in that population can be separated into the after-hyperpolarization (AHP) and a slow ramp leading to the next spike, as we have done in Fig. 4B. Previous modeling work achieved the slow rates of rise during the ramp was by carefully balancing the dynamics of K_V_4 and the inward L-type Ca_V_1.3 current (39). Here we showed that if the dynamics are constrained to a narrow region between the nullclines for voltage and inactivation of K_V_4, the balance between inward and outward currents does not need to be hand tuned to produce slow frequencies characteristic of convential DA neuron pacemaking. The nullcline for K_V_4 inactivation is simply the steady state inactivation curve as a function of membrane potential. The narrow region between the nullclines constrains the membrane potential to a ramp-like interspike interval that can be conceptualized as a slowly moving, pseudo “resting” potential. The evolution of the “resting” potential is driven by the time course of K_V_4 inactivation. An earlier study articulated the concept of a slowly moving fixed point (“resting” potential) but did not illustrate this concept (40). The key to the higher CVs exhibited by the atypical populations during pacemaking is their shallower AHP, which does not remove enough inactivation to effectively recruit K_V_4 that produces the ramplike ISI. The instantaneous IV curves in Fig. 5 reveal a much greater restorative slope conductance in the conventional (limited dynamic range) compared to atypical (extended dynamic range) models. Figure 4A shows a continuous barrage of current perturbations in vitro that are not reflected in deviations from the ramp-like voltage trajectory of the DMS projecting DA neuron, consistent with the strong restorative process we have described. Thus, we provide a mechanistic explanation for the higher CVs observed experimentally in the atypical populations compared to the conventional ones. Moreover, our results suggest that the 10% of the DA neurons that exhibited regular pacemaking activity *in vivo* (7), despite embedding within active circuits, were likely conventional DA neurons.

### Putative general principle for slow pacemaking

A recent influential review (41) on the mechanisms of pacemaking in mammalian neurons suggested that the mechanism for slow pacemaking in neurons is generally based on a steeply volt-age-dependent persistent sodium current with a midpoint near −60 mV. Specifically for slow pacemaking in regularly firing midbrain dopamine neurons, the current required during the interspike interval is extremely small, ∼−1 to −5 pA (29), thus the net ionic current during that period is approximately the size of the current carried by a single ion channel (41). Given the stochasticity of channel opening, it is extremely unlikely that a single channel is responsible. Instead, the review suggested that it is more likely that several macroscopic (both inward and outward) currents nearly cancel each other out during the ramp-like interspike interval, resulting in a net small inward current (42). Moreover, a previous computational study suggests that “achieving and maintaining a low firing rate is surprisingly difficult and fragile in a biological context (43)”. Thus, one commentary suggested that a small current carried by numerous membrane pores with a fS rather than a pS single channel conductance magnitude might be better suited to generate regular pacemaking (42). However, the mechanism that we suggest here with dynamics driven by a slow variable moving the trajectory through a channel near the voltage nullcline would remove the problem of precisely balancing the macroscopic inward and outward currents, favoring the balanced current hypothesis.

### Further Generalizion of this Mechanism

The dynamic mechanism for regularization of pacemaking by a slow variable described in this study may be a general biological organizing principle. For example, Fig. 1 of (44) shows that there is ramp phase in the pacemaking action potentials in the sinoatrial node of the rabbit. During this phase, as during the ramp phase of conventional, compressed firing range dopamine neurons, the net membrane current is very small suggesting the presence of sequential fixed points of the fast dynamics. As in this study, the sequence of “resting membrane potentials” may be driven by a slow variable. Moreover, acutely dissociated superchiasmatic neural pacemakers exhibited a highly regular spontaneous firing rate of 8.2 ± 3.8 Hz, with an average coefficient of variation of the interspike interval of 16 ± 10% (45). These cells also exhibit a ramp during the ISI. Another plausible generalization is to pedunculopontine cholinergic type II and noncholinergic type I neurons. The type II neurons fire rhythmically in vitro with a discharge rate from 3.5 to 16.5 Hz and the ISI also has a ramplike appearance. The ISI ramp is likely due to an A-type current as in the DMS projecting dopamine neurons in study, because they also display a ramp response after a hyperpolarizing step due to removal of inactivation from an A-type potassium current (Figure S2B). Tellingly, the noncholingergic (type I) neurons also fire spontaneously, but they lack an A-type current and their firing is irregular rather than rhythmic (46). Note that the less regular pacemaking population in our study, the mNAcc projecting VTA neurons, possess an A-type K+ current that can be evoked by a hyperpolaring ramp (supplementary figure S2A), but that current is not recruited during pacemaking. A final candidate for generalization of the mechanism for robustness is the tonically active cholinergic neurons (TANs) in the striatum. The presence of a spontaneous baseline firing rate so that bidirectional changes in firing are possible, as in dopamine neurons, is likely related to their reward related signaling properties. Whereas dopamine neurons decrease their firing to a worse than expected reward, TANs learn to decrease their firing in response to a stimulus that predicts a reward (47). In vitro, they exhibit a spontaneous regular firing (rate, 2.45–4.03 Hz; CV, 0.10–0.19). These cells also exhibit a depolarizing ramp to spike threshold during spontaneous tonic firing. Moreover, when firing rate was decreased by a hyperpolarizing step current, the CV increased linearly as the firing rate decreased (48), somewhat similarly to our model neurons (Supplementary Figure 3).

### Basal Ganglia Pacemakers

Most principal cells in the basal ganglia are pacemakers, with the lone exception of the medium spiny striatal cells (3). In the three examples given in this section, block of synaptic transmission did not affect rate or regularity of spontaneous pacemaking observed in vitro. For example, glutamatergic subthalamic nucleus (STN) neurons in rat slices fired rhythmically at room (3.6 Hz with 0.12 CV) or body temperature (6.5 Hz with 0.11 CV) (49). The shape of the ISI was different in the STN compared to conventional DA cells. In our model and in many neuromodulatory cells described above, the depolarizing phase of the ISI has two slopes: the first one is attributed to SK turning off and ending the AHP, whereas the second one is the nearly constant slope of the ramp phase. In STN cells (Fig 1C, 4AC of (49)), the initial slope is either flat or nonexistent, suggesting that perhaps the SK current itself is the operant slow process. This is supported by the long duration of the hyperpolarizing phase of the AHP in those cells. However, in the presence of Cd or an SK blocker, regularity was rescued at faster than physiological firing rates, suggesting a slow process other than the SK current that is activated by depolarization, possibly similar to the TANS. In fast-firing dissociated cells, slow inactivation of the sodium channels appears to be the dominant slow process (50). The intrinsic ability of STN neurons to fire rhythmically likely allows those cells to provide a fairly constant background excitatory tone to basal ganglia structures in the absence of any specific inputs.

GABAergic globus pallidus (GP) neurons also exhibited single-spike pacemaker activity, at 12.5±0.4 Hz with a CV of 0.18±0.01 in mouse slices (51). In these neurons, the hyperpolarizing phase of the AHP was quite brief and most of the ISI consisted of a depolarizing ramp. The HCN channel blocker ZD7288 significantly slowed the discharge rate and decreased regularity, suggesting that the slow inactivation of this current might provide a slow variable to regularize the firing, and its activation variable may provide the restorative force. In contrast to the TANs, however, there was no correlation between CV and discharge rate. The authors of this study theorized that one role of pacemaking in this population might be to create a synchronizing mechanism via phase resetting of the pacemaker response by striatal GABAergic input.

Similarly, GABAergic substantia nigra pars reticulata (SNr) neurons in rat slices fired at 11.35 ± 3.27 Hz with a CV of 0.044 ± 0.022 (52). The rate and regularity were unchanged by block of the HCN channel. The ISI waveform appeared more similar to the STN neurons rather than the globus pallidus neurons. SK block shortened the duration of the AHP by eliminating the slower component of the AHP while leaving the fast component intact, again similar to STN neurons. This resulted in a tripling of the CV but the Cv is apamin was still ∼0.15, suggesting the possible presence of another regularizing influence,

### Summary

Our results suggest that in order for pacemaking to be robust, at least one slow variable relative to the spiking currents is required to create an effective and slowing changing “resting potential”. In our example of regular pacemakers, the DMS-projecting conventional SNC neurons, the slow variable is inactivation of the A-type potassium current, in this case mostly mediated by K_V_4.3. In addition, a fast component of a restorative current is required to oppose fluctuations from this effective resting potential. In our example, the relatively fast activation and de-activation of the same current provides both the slow variable and the restorative force, but this need not be the case in other neurons. As there are many examples of pacemaking neurons in the mammalian central nervous system, some may share this A-type K^+^ current mechanism, but other slow variables such as inactivation of the persistent sodium current, inactivation of the H current, and Ca^2+^ activation of the SK channel may be the determinants of regularity in other populations. There are many situations in which a background level of neurotransmission in the target area is advantageous, thus the mechanisms we present here have general applicability.

### Computational Methods

#### Computational Model Development

Models of a representative neuron in each of the three DA subpopulations were created using the same currents shown in the equivalent circuit diagram in Fig. 2, with differences in maximal conductances given in Table 1. One compartment models were chosen because, as we demonstrate below, they are amenable to mathematical analysis in the phase plane. The reduction of dimensionality required for this analysis is not feasible for a heterogeneous multicompartment model.

We and others (16,53,54) have previously successfully used single compartment models to capture the dynamics of experimentally observed phenoma. In addition to the equivalent circuit, the cytosolic Ca^2+^ dynamics were taken into consideration to model the differential engagement of SK channels in the DA subpopulations. We assumed that the pool of Ca^2+^ sensed by the SK channel was a microdomain fed by nearby Ca^2+^ channels. Due to the limitations of the NEURON software package, the SK microdomain was implemented as a thin shell for convenience – with calcium extrusion via the pump occurring from the nominally ‘interior’ shell representing the remainder of the cytosol. A true microdomain only senses Ca^2+^ from Ca^2+^ channels that are within a few microns. The thin outer shell coupled to a larger Ca^2+^ pool is a phenomenological model to capture the dynamics that drive the SK channel rather than a representation of the time course of bulk Ca^2+^. The fractional Ca^2+^ coupling to each Ca^2+^ channel was based on experimental observations that the SK channel is primarily coupled to high threshold Ca^2+^ channels (represented by N-type Ca^2+^ channels in this model) in order to produce the AHP (24), with a contribution from T-type (55) but with limited coupling to the L-type (56) over pacing timescales. Both microdomain and bulk compartments incorporate dynamic buffering via binding and unbinding through a mass balance with a single buffer (57,58). Population level differences in the calcium binding proteins (59) were neglected, thus the buffering capacity and rates of calcium binding were assumed to be constant across subpopulations. The shallower AHP amplitude in the atypical, limited dynamic range subpopulation is the single most reliable functional property for that distinguishes that subpopulation (9,60). Accordingly, the model for the VTA-mNAcc projecting DA cells has a 3-5 fold lower maximum SK conductance and as well a larger volume for Ca^2+^ accumulation that decreases the maximum concentration of Ca^2+^ seen by each SK channel and the maximum current evoked by a given Ca^2+^ influx. This corresponds conceptually to a hypothesized larger mean separation between SK and Ca^2+^ channels. Whereas conventional neuron exhibit a subthreshold oscillation during pacemaking (61,62), the atypical neurons located in the VTA lack a subthreshold oscillation Ca^2+^ to recruit SK during pacemaking (63,64), thus the Ca2+ conductances for the L and T-type conductances were also reduced in the medial shell projecting model.

The fast sodium channel responsible for action potential upstroke is implemented in a Markov model (16) that includes a long-term inactivation state. In this study the longterm inactivated state was included only in the conventional DMS projecting population. Action potential repolarization is achieved with a delayed rectifying potassium channel that qualitatively includes the effects of multiple fast activating potassium channels, such as BK and fast components of Kv4.3 that its HH formulation cannot adequately capture. The AHP is generated by a combination of SK, adapted from (39), and KM (Kv7) adapted from (20). The K_V_4.3 channel was adapted from (21), with KChip-subunits mediating fast (∼50 ms) and slow (∼200 ms) inactivation (65,66) in the mNAcc projecting population. The DMS-projecting models assume that only KChip-subunits mediating fast inactivation are present. The relative values of AHP currents were chosen to replicate the effects of apamin (24) on AHP depth. The ERG channel (Kv11) (67) was included in all DA subtypes to regulate pacemaker frequency, particularly under simulated apamin application, and to allow for consistent repolarization from depolarized states under simulated spiking blockers. The K-ATP channel (68) was neglected since coupling between metabolic and electrical activity is not essential for the phenomena we model here. The N and L-type Ca^2+^ currents are included in all populations, with the N-type channel (CaV2) representing a composite of all high-threshold calcium channels (N, P/Q, R). The H current (HCN) (69)is calibrated to produce an electrophyisio-logically relevant sag potential in the conventional DMS-projecting model population and omitted from the atypical medial shell projecting population. The model was developed in python and NEURON (70) and is freely available on Modeldb at https://modeldb.science/2041613 (reviewer password: nullcline). Additionally, full model equations can be found in the supplemental materials.

### Experimental Methods

All experimental procedures involving mice were approved by the German Regional Council of Darmstadt (V54-19c20/15-FU/1257).

#### Methods - In Vitro Patch, Retrograde Tracing

C57BL/6N mice were anesthetized using isoflurane (AbbVie, USA; induction, 3.5%; maintenance, 0.8 to 1.4% in O2, 0.35 L/min) and placed in a stereotaxic frame (Kopf). A topical anaesthetic (lidocaine gel; EMLA crème, AstraZeneca, UK) was applied to the incision site. Throughout surgery, body temperature, respiratory rate (1–2 Hz), and reflexes were continuously monitored. Craniotomies were performed using a stereotaxic drill (0.5 mm diameter) to mNAcc (bregma: 1.54 mm, lateral: ±0.45 mm, ventral: −4.1 mm), lNAcc (bregma: 0.86 mm, lateral: ±1.75 mm, ventral: −4.5 mm), DLS + (bregma:, +0.74 mm; ML, 2.2 mm; DV, 2.6 mm), and DMS (bregma: +0.74 mm; ML, 1.2 mm; DV, 2.6 mm). Red retrobeads (100nl; Lumaflor) diluted (1:30) in artificial cerebrospinal fluid (ACSF; Harvard Apparatus) were injected either in the mNAc or lNAc using a 1 μL Hamilton syringe (Hamilton, Switzerland). Patch-clamp experiments were performed 2 to 4 days after tracer injection.

#### Slice preparation

Mice were anesthetized by intraperitoneal injection of ketamine (250 mg/kg; Ketaset, Zoetis) and medetomidine hydrochloride (2.5 mg/kg; Domitor, OrionPharma) before intracardial perfusion using ice-cold ACSF (containing 125 mM NaCl, 2.5 mM KCl, 6 mM MgCl2, 0.1 mM CaCl2, 25 mM NaHCO3, 1.25 mM NaH2PO4, 50 mM sucrose, 2.5 mM glucose, 3 mM kynurenic acid, oxygenated with 95% O2 and 5% CO2). The brain was harvested and the midbrain was cut into 250 μm thick coronal slices using a vibratome (Leica VT1200S, Leica Biosystems, Germany). Before the experiment, slices recovered for 1h at 37°C in oxygenated ACSF (125 mM NaCl, 3.5 mM KCl, 1.2 mM MgCl2, 1.2 mM CaCl2, 25 mM NaHCO3, 1.25 mM NaH2PO4, 22.5 mM sucrose and 2.5 mM glucose).

#### In vitro electrophysiology

For patch-clamp experiments, the slices were transferred to a temperature-controlled recording chamber maintained at 37 °C (Temperature Controller VI, Luigs & Neumann, Germany) and consistently perfused with ACSF at a rate of 2–4 ml/min. To block synaptic transmission, CNQX (20 μM, Biotrend), DL-AP5 (10 μM, Tocris), (-)-Sulpiride (0.15µM, Tocris), CGP (50 nM, Tocris) and Gabazine (SR95531 4 μM, Biotrend) were added to the ACSF.

Neurons were visualized using a light microscope (Axioskope 2 FS plus, Zeiss, Germany) equipped with an infrared light-source (SOLIS-850C, ThorLabs, USA) and a digital camera (CS505MUP1, ThorLabs, USA). Retrogradely traced neurons were identified by excitation of red retrobeads with a filtered LED lamp (SOLIS-1C, ThorLabs, USA, filtered at 546/12 nm). Labeled DA Neurons were recorded using borosilicate glass pipettes (3–4 MΩ, GC150TF, Harvard Apparatus, USA) filled with an internal solution (135 mM K-gluconate, 5 mM KCl, 10 mM HEPES, 0.1 mM EGTA, 5 mM MgCl2, 0.075 mM CaCl2, 5 mM ATP, 1 mM GTP, 0.1% Neurobiotin, pH 7.35, 290–300 mOsmol). Recordings were obtained using an EPC-10 patch-clamp amplifier (HEKA Elektronik) at a sampling rate of 20 kHz and a low-pass Bessel filter (5 kHz).

Data acquisition and analysis were performed using PatchmasterNext (HEKA Elektronik, Germany) and MATLAB (MathWorks, USA), and statistical analyses were conducted with GraphPad Prism 10 (GraphPad Software). Only spontaneously active midbrain dopaminergic neurons showing stable pacemaker firing were included for analysis. Neurobiotin-filled neurons were verified post hoc by immunohistochemistry.

#### Immunohistochemistry

Midbrain slices and harvested forebrains were stored overnight at 4 °C in a fixative solution consisting of 4% paraformaldehyde and 0.15% picric acid in phosphate-buffered saline (PBS; pH 7.4). On the following day, the tissue was transferred to a storage solution containing 10% sucrose and 0.05% NaN₃ in distilled water, and stored at 4 °C. Forebrains containing the striatum were cut into 80 µm-thick coronal slices using a vibrating microtome (VT1000S, Leica Biosystems, Germany). Slices were washed in PBS (0.2 M, pH 7.4) and then incubated in a blocking solution (0.2 M PBS containing 10% horse serum, 0.5% Triton X-100, and 0.2% BSA) for 1 h for striatal slices and 2 h for thicker patch slices. Subsequently, slices were incubated overnight at room temperature in a carrier solution (0.2 M PBS containing 1% horse serum, 0.5% Triton X-100, and 0.2% BSA) with anti-tyrosine hydroxylase antibody (1:1000; Millipore, Germany). The following day, slices were washed in PBS and incubated overnight at room temperature in carrier solution containing goat anti-rabbit Alexa Fluor 488 (1:750; Invitrogen, USA) and streptavidin–Alexa Fluor 568 (1:750; Invitrogen, USA). On the third day, slices were washed in PBS, incubated in DAPI (0.2 µL/mL) for 5 min, mounted on glass slides using Vectashield mounting medium (Vector Laboratories, USA), and stored at 4 °C.

---

**Figure S1:**
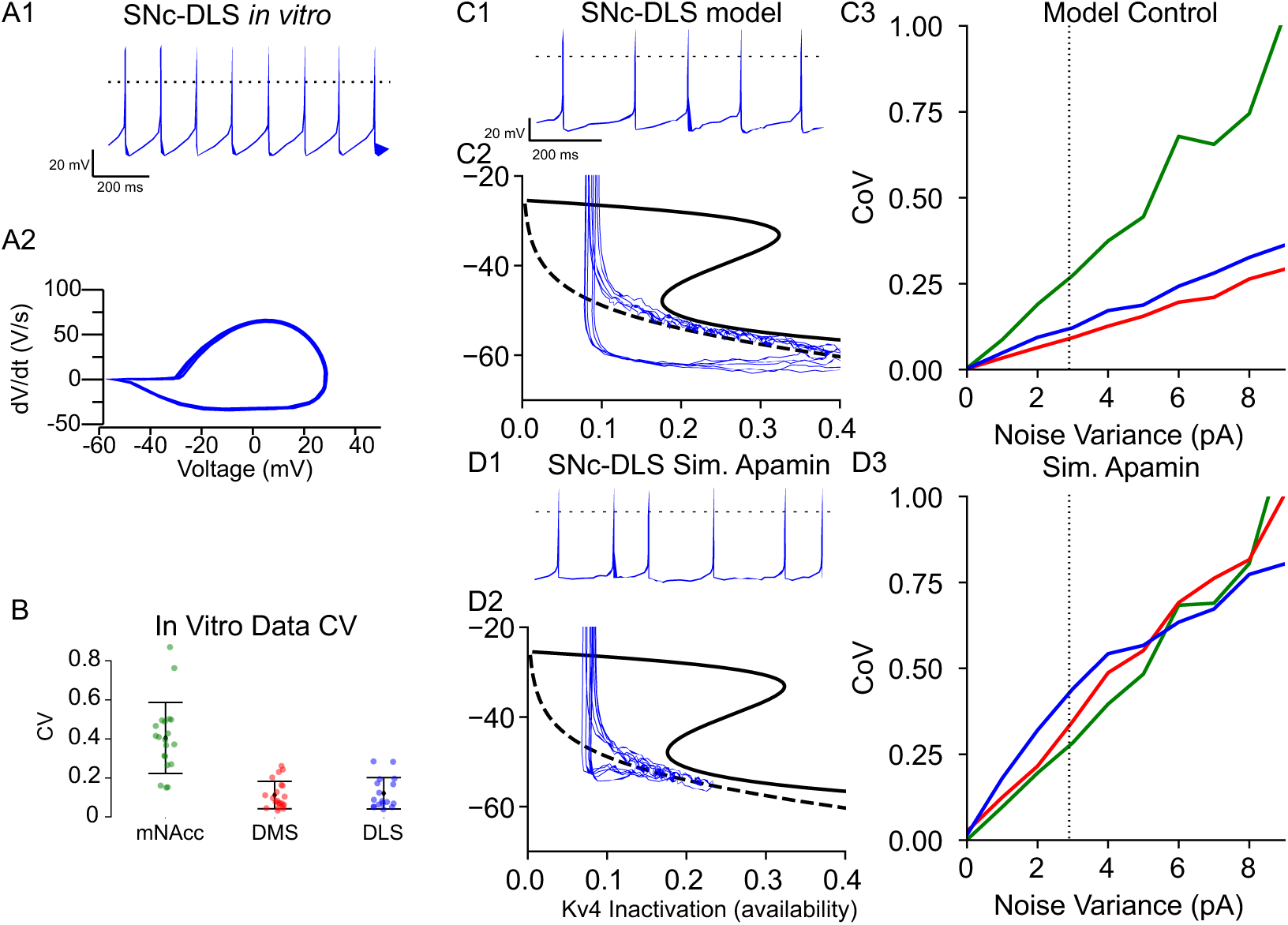
DLS projecting cells mirror those of DMS in CV and in response to simulated apamin. A. Representative in vitro trace of identified DLS projecting cell in vitro. A1 Voltage vs time. A2 dV/dt vs V for A1. B. CV data for N=20 DLS projecting cells. mNAcc, DMS reproduced from Figure 6 for reference. C,D: DLS projecting model under simulated in-vitro, simulated apamin as in Fig. 5. DLS CV blue with DMS (red), and mNAcc (green) reproduced from Fig 5. Model parameters are as those for SNc-DMS with the following changes: gNaL=4.5, gCaT=30, tauKA=35 ms, gSK=150, gKv4=281.25 (all uS/cm^2^ unless indicated)

**Figure S2:**
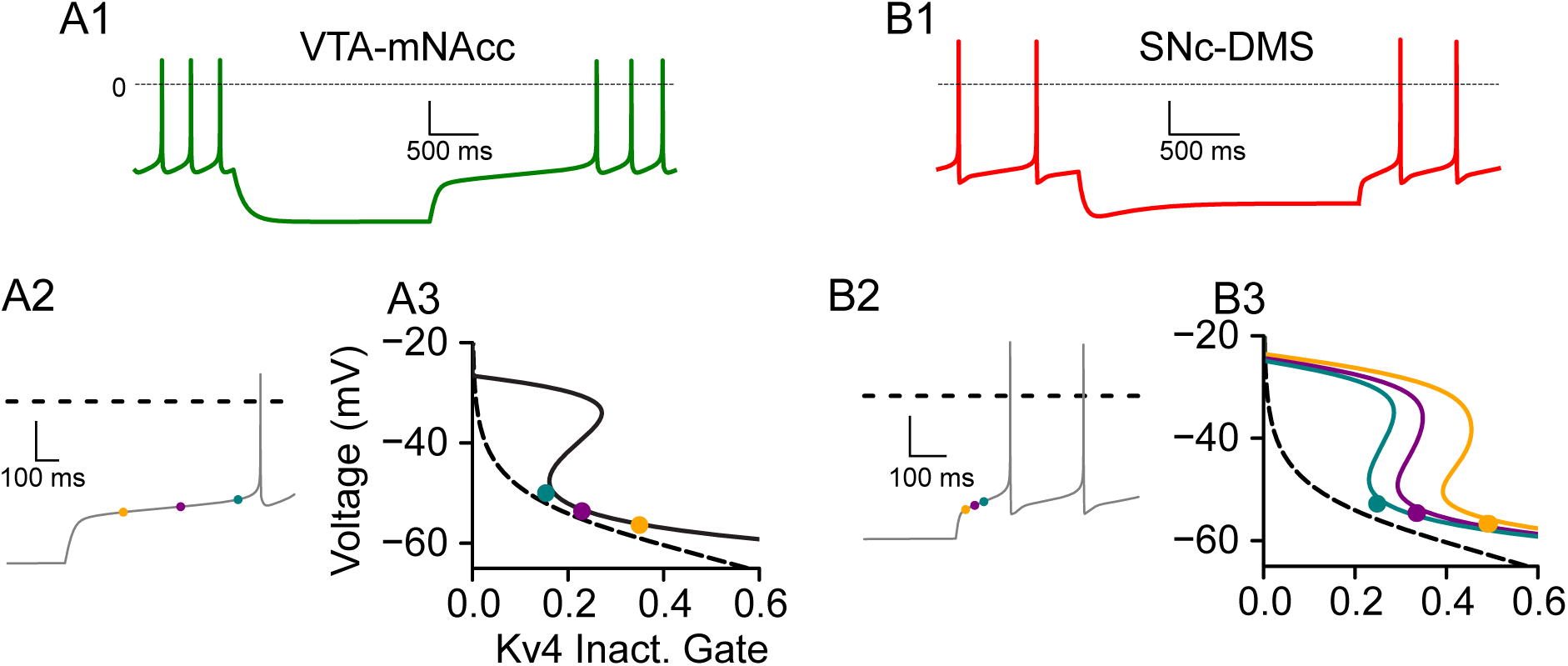
Simulated ramp rebound responses in mNAcc and DMS projecting models. A. VTA-mNAcc model exhibits long (2s) ramp following recovery from hyperpolarization to −80 mV. A1: full trace, A2 zoom in of rebound, A3: Kv4 inactivation gate, voltage phase space as in Fig 4. While Kv4 is not significantly recruited during pacing, it contributes strongly to post-inhibitory rebound responses. B. SNc-DMS as in A. Hyperpolarization of SNc-DMS model to −80 mV shows characteristic ‘sag’ potential produced by the H current, 300 ms rebound ramp. Despite presence of inward hyperpolarization activated currents (I_Ca,T_, I_H_), the A-type current dominates during the ramp upon rebound from hyperpolarization in this population. Colored nullclines in B3 correspond to voltage nullclines with the slow dynamics of inward hyperpolarization set to their instantaneous values at the indicated points.

**Figure S3:**
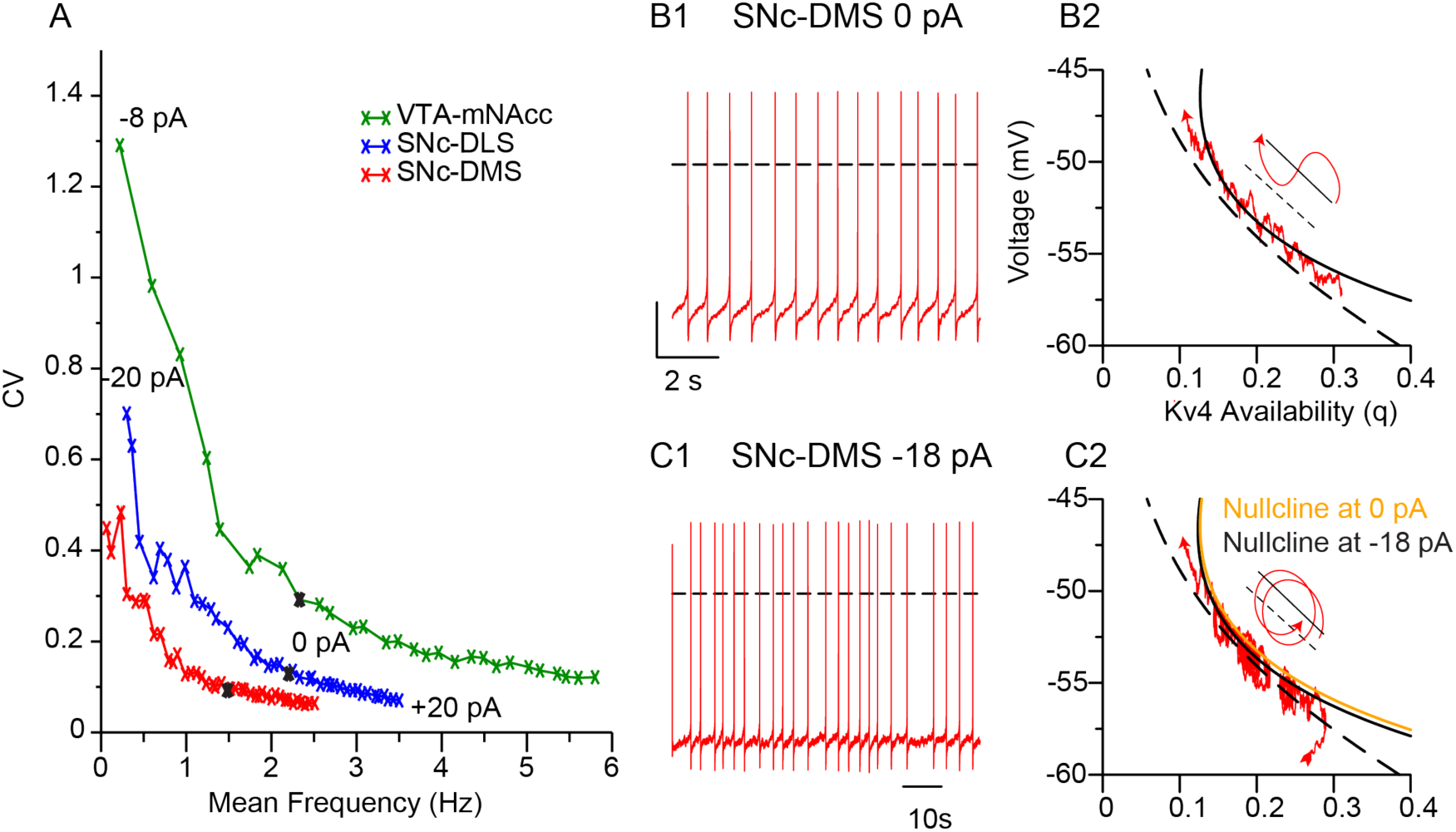
Variability increases with decreasing firing rate. A. CV vs Mean Frequency for each model (VTA-mNAcc: green, SNc-DMS: red, SNc-DLS blue) as DC current is applied from −20 pA to 20 pA in intervals of 1 pA under simulated in vitro channel noise. Lower frequencies have larger CV for all populations. Black crosses indicate 0 pA. B. DMS projecting model under simulated channel noise and 0 applied current (Same model as Fig 5A). B1 Voltage vs Time. B2 Zoom in to ramp region of Kv4 phase space from Fig 5A1. Noise induce perturbations do not cross Kv4 nullcline (dashed line). Trajectories oscillate about voltage nullcline, but Kv4 availability (q) monotonically decreases. C. Same as B but on a much slower time scale (compare scale bars) for −18 pA DC current. Regularity is lost at very low frequencies as Kv4 inactivation is no longer sufficiently slow compared to voltage. At sufficiently low frequencies, the random walk evoked by simulated channel noise can become large relative to the mean net current. During the ramp region, perturbations routinely cross Kv4 nullcline, removing Kv4 inactivation. Noise driven crossings result in spiraling dynamics with a slow drift to top left.

